# The parotid secretory protein BPIFA2 is a salivary surfactant that affects LPS action

**DOI:** 10.1101/2020.02.08.940163

**Authors:** Seshagiri R. Nandula, Ian Huxford, Thomas T. Wheeler, Conrado Aparicio, Sven-Ulrik Gorr

**Author notes:** Corresponding author: Sven-Ulrik Gorr.

## Abstract

Saliva plays important roles in the mastication, swallowing and digestion of food, speech and lubrication of oral mucosa, antimicrobial and anti-inflammatory activity and control of body temperature in grooming animals. The salivary protein BPIFA2 (BPI fold containing family A member 2; former names: Parotid Secretory Protein, PSP, SPLUNC2, C20orf70) is related to lipid-binding and LPS-binding proteins expressed in mucosa. Indeed, BPIFA2 binds LPS but the physiological role of BPIFA2 remains to be determined. To address this question, *Bpifa2* knockout (Bpifa2^tm1(KOMP)Vlcg^) (KO) mice were phenotyped with a special emphasis on saliva and salivary glands. Saliva collected from KO mice was less able to spread on a hydrophobic surface than wild-type saliva and the surface tension of KO saliva was close to that of water. These data suggest that BPIFA2 is a salivary surfactant that is mainly responsible for the low surface tension of mouse saliva. The reduced surfactant activity of KO saliva did not affect consumption of dry food or grooming, but saliva from KO mice contained less LPS than wild-type saliva. Indeed, mice lacking BPIFA2 responded to ingested LPS with an increased stool frequency, suggesting that BPIFA2 plays a role in the solubilization and activity of ingested LPS. Consistent with these findings, BPIFA2-depleted mice also showed increased insulin secretion and metabolomic changes that were consistent with a mild endotoxemia. These results support the distal physiological function of a salivary protein and reinforce the connection between oral biology and systemic disease.

## Introduction

Saliva plays important roles in the mastication, swallowing and digestion of food, speech and lubrication of oral mucosa, antimicrobial and anti-inflammatory activity (Amerongen and Veerman, 2002; Dawes et al., 2015; Gorr, 2009) and control of body temperature in grooming animals (Obal et al., 1979). While well over 2000 proteins have been identified in human saliva (Loo et al., 2010), the physiological function of several abundant proteins remains poorly defined.

BPIFA2 (BPI fold containing family A member 2 (Bingle et al., 2011); former names: Parotid Secretory Protein (PSP), SPLUNC2, C20orf70) was first described as a leucine-rich protein in salivary glands (Wallach et al., 1975) and identified as Parotid Secretory Protein in mouse parotid glands (Owerbach and Hjorth, 1980). BPIFA2 is abundantly expressed in all three major mouse salivary glands (Gao et al., 2018) and human salivary glands (Geetha et al., 2003). The protein has been isolated from human saliva (Geetha et al., 2003) where it represents about 5% of total protein (Ruhl, 2012).

We cloned the cDNA for human BPIFA2 (PSP) (Venkatesh et al., 2001) and noted that the translated amino acid sequence was predicted to form a similar protein fold to that of bactericidal/permeability-increasing protein (BPI) and LPS-binding protein (LBP) (Geetha et al., 2003). Several related genes have been identified on human chromosome 20q11 (Bingle and Craven, 2004; Bingle and Gorr, 2004; Weston et al., 1999). The corresponding proteins are predominantly expressed in mucosal tissues and include the airway protein PLUNC (BPIFA1) (Weston et al., 1999), which binds LPS (Ghafouri et al., 2004) and acts as a surfactant with anti-biofilm activity in airway epithelia (Gakhar et al., 2010). Another noted family member is latherin (BPIFA4), a surfactant in horse sweat and saliva (Beeley et al., 1986; McDonald et al., 2009).

A number of possible functions and physiological effects have been suggested for BPIFA2, including cleavage in Sjögren’s syndrome (Robinson et al., 1998a) and lipid binding and protein transport in secretory granules (Venkatesh et al., 2011). BPIFA2 binds candida (Holmes et al., 2014), LPS (Abdolhosseini et al., 2012), and bacteria (Robinson et al., 1997). The latter presumably explains the observed bacterial agglutination at relatively high concentrations of BPIFA2 (Gorr et al., 2011). A proposed bactericidal role for BPIFA2 remains controversial (Geetha et al., 2003; Gorr et al., 2011; Prokopovic et al., 2014). Thus, the physiological role of BPIFA2 in saliva remains to be determined.

To determine further the physiological role of BPIFA2, we have analyzed a mouse knockout model (*Bpifa2* KO). *Bpifa2* KO mice have recently been phenotyped and the results posted by the International Mouse Phenotyping Consortium (IMPC) (Brown et al., 2018). This effort analyzed a variety of growth, developmental and metabolic parameters without identifying an obvious phenotype for the BPIFA2 deficient mice (IMPC, 2019). However, salivary glands and saliva, the primary locations of BPIFA2 expression were not included in this analysis. To address this gap, we analyzed Bpifa2^tm1(KOMP)Vlcg^ mice with a special emphasis on saliva and salivary glands.

## Materials and Methods

### Generation of *Bpifa2* knock-out mice

All animal experiments were reviewed and approved by the Institutional Animal Care and Use Committee of the University of Minnesota. *Bpifa2* knockout mice (Bpifa2^tm1(KOMP)Vlcg^) (IMSR Cat# KOMP:VG14843-1-Vlcg, RRID:IMSR_KOMP:VG14843-1-Vlcg) were generated on a C57BL/6NTac background using ES colonies that were generated by Regeneron Pharmaceuticals, Inc. using the approach described by Regeneron (http://velocigene.com/komp/detail/14843) (Valenzuela et al., 2003). ES cell injection and strain validation was conducted by the KOMP at UC Davis (https://www.komp.org), which delivered heterozygous and wild-type breeder mice to our laboratory in June, 2012. The mice were housed at the University of Minnesota under specific pathogen free (SPF) conditions with free access to water and irradiated rodent chow (Envigo (Harlan) Teklad 2918), unless otherwise noted for individual experiments. Mice were initially bred as heterozygous crosses. As viability was established, homozygous crosses were used. Genotyping of tail-snips from mice resulting from heterozygous crosses was performed at age 3 weeks by the University of Minnesota Genomics Center using the primers listed in **Table 1** or by Transnetyx (Cordova, TN) using real-time PCR with specific probes designed for *Bpifa2*. KO and WT mice were from the same colony, of similar age (except as noted) and housed under identical conditions.

**Table 1:**
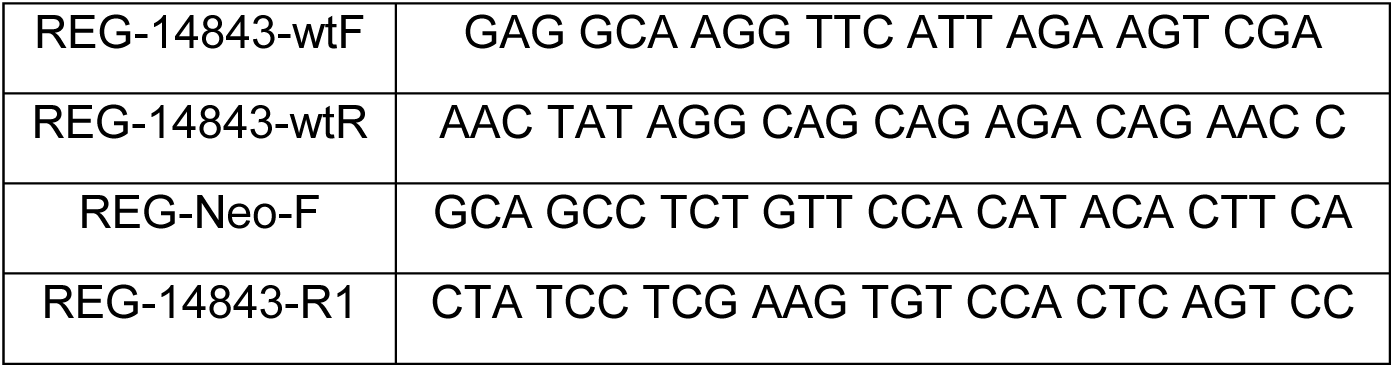
PCR primers for mouse genotyping.

### Saliva collection

Mice were fasted for 2-3h, weighed and anesthetized with ketamine (75 mg/kg) and xylazine (8 mg/kg) (i.p.) and then injected s.q. with pilocarpine (Sigma-Aldrich) (0.5 mg/kg body weight) that was freshly prepared in dH_2_O (75 µg/ml). Saliva was collected for 20 min, using a micropipette, and immediately stored on ice. The volume was determined with a micropipette. Saliva samples were centrifuged for 4 min at 2,000 x g and the supernatants stored at −20°C until use.

### SDS-PAGE and immunoblotting

Saliva supernatant was analyzed by SDS-PAGE and stained with Coomassie G-250 (Simply Blue; Thermo Fisher). Relative molecular weight estimates were based on the migration of unstained molecular weight markers (Precision Plus Protein; BioRad, Hercules, CA).

Proteins separated by SDS-PAGE were also transferred to PVDF membranes, which were blocked with 2% Tween 20 in Tris-buffered saline (TBS) for 3 min and then washed in TBS (3×10 min). The membranes were then incubated with an affinity-purified goat-anti-mouse BPIFA2 antibody (EB10621) (Everest Biotech, Upper Heyford, UK) (diluted 1:5000 in TBS) overnight at room temperature. The blots were again washed and incubated with HRP-conjugated rabbit anti-goat IgG (H+L) secondary antibody (Invitrogen) (diluted 1:10,000 in TBS) for 45 min at room temperature. After final washes, the membranes were incubated with ECL Western blotting substrate (Thermo Scientific Pierce, Rockford, IL) and chemiluminescence visualized in a Kodak Image Station 4000R.

### Surface tension and water contact angles of mouse saliva

The surface tension of mouse saliva was determined with the pendant drop method using the DM CE-1 contact angle meter (Kyowa Interface Science Co. Ltd., Saitama, Japan). Saliva drops of 5-6 µl were formed at the tip of a syringe and the shape of the pendant drop analyzed using the FAMAS software (Kyowa Interface Science Co. Ltd., Saitama, Japan). The experiment was performed in triplicate.

To prepare hydrophobic substrates, silica microscope cover slides were sonicated for 10 min sequentially in cyclohexane, deionized water, and acetone, and finally dried with a stream of N_2_ gas. After cleaning, the slides were stored in a glass petri dish cleaned with 2% Hellmanex solution (Hellma Analytics, Müllheim, Germany) under vacuum until surface modification. For hydrophobic surface modification, cleaned slides were placed in Erlenmeyer flasks and flooded with N_2_. Ten ml silanization solution (0.4% dichloromethylsilane in trichloroethylene) (Sigma-Aldrich, MO, USA) was injected into the flask. The slides in solution were sonicated for 2 min, and left for 10 min. This was repeated five times, 60 min total. Samples were removed from the flasks with forceps that were previously cleaned with with 2% Hellmanex solution. Slides were rinsed sequentially in cleaning solutions of ethanol, isopropyl alcohol, deionized water, and acetone followed by drying with a stream of N_2_. Slides were placed in a vacuum desiccator for 24 hours after surface treatment to further dry the silanized surface.

Dynamic contact angles between mouse saliva and the hydrophobic substrate were determined with the sessile drop method using the DM CE-1 contact angle meter. Two µl drops of mouse saliva supernatant were dispensed on silanized glass slides (hydrophobic surface) and the contact angles measured using the FAMAS software. Contact angles were measured in duplicate for two different mice per group and the wettability data expressed as the decrease in contact angle from time 0s to 20s.

### Food and water consumption

Mice were weighed and housed singly or in pairs with free access to powdered rodent chow for one week prior to quantitation of food consumption. Food dishes were weighed before and after re-filling at 1-3 day intervals. The average consumption from 3-6 determinations for each cage was expressed as g food/day/kg body weight.

To determine water consumption, mice were housed singly or in pairs with free access to food and water for a week. Water bottles were weighed before and after refilling and the daily consumption per mouse calculated for each group on 4-5 consecutive days.

### Histology

Submandibular glands were collected immediately after euthanasia, fixed in buffered formalin and embedded in paraffin. Paraffin sections were cut and H&E stained by a commercial laboratory (Histoserv, Germantown, MD). Tissue sections were viewed by light microscopy at 50-400x magnification. The investigator was blinded to the genotype and sex of the animals at the time of examination. Inflammatory infiltrates (foci) consisting of at least 50 cells were identified by their blue stain in H&E stain and counted in each tissue section.

### LPS assays

Blood was collected into EDTA-tubes from mice that had been fasted for 12h. LPS activity was detected in saliva or blood plasma using hTLR4-HEK reporter cells (Invivogen, San Diego,CA). Samples (20 µl) were mixed with 180 µl cell suspension in HEK-Blue detection medium (Invivogen) and plated in 96-well plates (25,000 cells/well). The cells were cultured overnight at 37°C, humidified atmosphere with 5% CO_2_. LPS activity was determined by the induction of secreted alkaline phosphatase, which was determined spectrophotometrically at 630 nm, as described by the supplier (Invivogen). The samples in each independent experiment were normalized to the mean of the WT samples in that experiment. Saliva samples were also tested by a chromogenic Limulus Amebocyte Assay (Thermo Scientific Pierce, Rockford, IL), which yielded qualitatively similar results (not shown).

### Stool collection

Groups of 2 or 4 mice were maintained without water for 12 h and then provided drinking water with or without 1 mg/ml LPS. Mice were acclimatized for 4-6 hours. Cages were then cleaned and feces was counted and weighed at the end of a 2 or 18 hour collection period. Neither protocol resulted in diarrhea in LPS-fed or control mice. The two protocols yielded similar results and the data were combined and expressed as fecal pellets/mouse/hour or mg feces/mouse/hour.

### Glucose determination

Mice were fasted for 2 hours and glucose was determined in freshly collected blood from groups of four mice aged 6 months, using a commercial glucometer (FreeStyle, Abbott Diabetes Care, Alameda, CA or True Result, Nipro Diagnostics, Ft. Lauderdale, FL). The same mice were re-analyzed at ages 8-9 months.

### Insulin ELISA

Mouse insulin was determined by ELISA (Crystal Chem Cat# 90080, RRID:AB_2783626) following the manufacturer’s instructions (Crystal Chem, Elk Grove Village, IL). Blood was collected from animals aged 3-6 months that were without food for <2 hours before blood collection. Either serum or plasma was prepared and stored at −20C until analysis.

### Metabolomic analysis

Targeted metabolomics was conducted with the Biocrates Absolute IDQ p180 kit (BIOCRATES Life Sciences AG, Innsbruck, Austria) by the West Coast Metabolomics Center, UC Davis, (Davis, CA). Blood plasma from fasted male and female, WT and KO mice (10 male and 8 female mice per group) was stored on dry ice and analyzed within 3 months (Wagner-Golbs et al., 2019). Ten µl aliquots were used for the analysis of five groups of metabolites, i.e. acylcarnitines, amino acids, biogenic amines, glycerophospholipids and sphingolipids. The average metabolite concentration in each group (N=8-10) was calculated and the ratio KO/WT for each metabolite was determined for each sex. The P-values for KO and WT samples of each sex were calculated by unpaired, two-tailed Student’s test. Volcano plots were used to show the P-values as a function of the KO/WT ratio for each metabolite.

## Results

BPIFA2 is expressed in human saliva (Geetha et al., 2003) and an analysis of BPIFA2 expression in 44 human tissues confirmed the salivary gland specific expression of this protein (proteinatlas.org). Consistent with these findings, an analysis of 66 tissues in *Bpifa2* KO mice show *Bpifa2* gene expression is limited to salivary glands and the tongue with smaller amounts detected in penis tissue (kompphenotype.org). Indeed, the *Bpifa2* (*Psp*) promotor has been utilized to achieve salivary gland specific expression of a transgene, with negligible expression detected in other parts of the GI tract (Golovan et al., 2001).

Analysis of WT and KO mouse saliva by SDS-PAGE (Fig. 1A) and immunoblotting (Fig. 1B), revealed a strong band at 22kD was seen in WT mice but not in KO mice (Fig. 1A). Despite some individual differences, no other consistent changes in protein expression were determined in a comparison of three mice from each group. The migration of the 22kD band is consistent with that of mouse BPIFA2 (PSP) (Owerbach and Hjorth, 1980), which was confirmed by immunoblotting with an antibody to mouse BPIFA2 (Fig. 1B). The main BPIFA2 band and a smaller, presumable, degradation product were visible in saliva from WT mice. The bands were absent in saliva from KO mice and the BPIFA2 bands were detected at a reduced intensity in saliva from heterozygous mice (Fig. 1B).

**Fig. 1.**
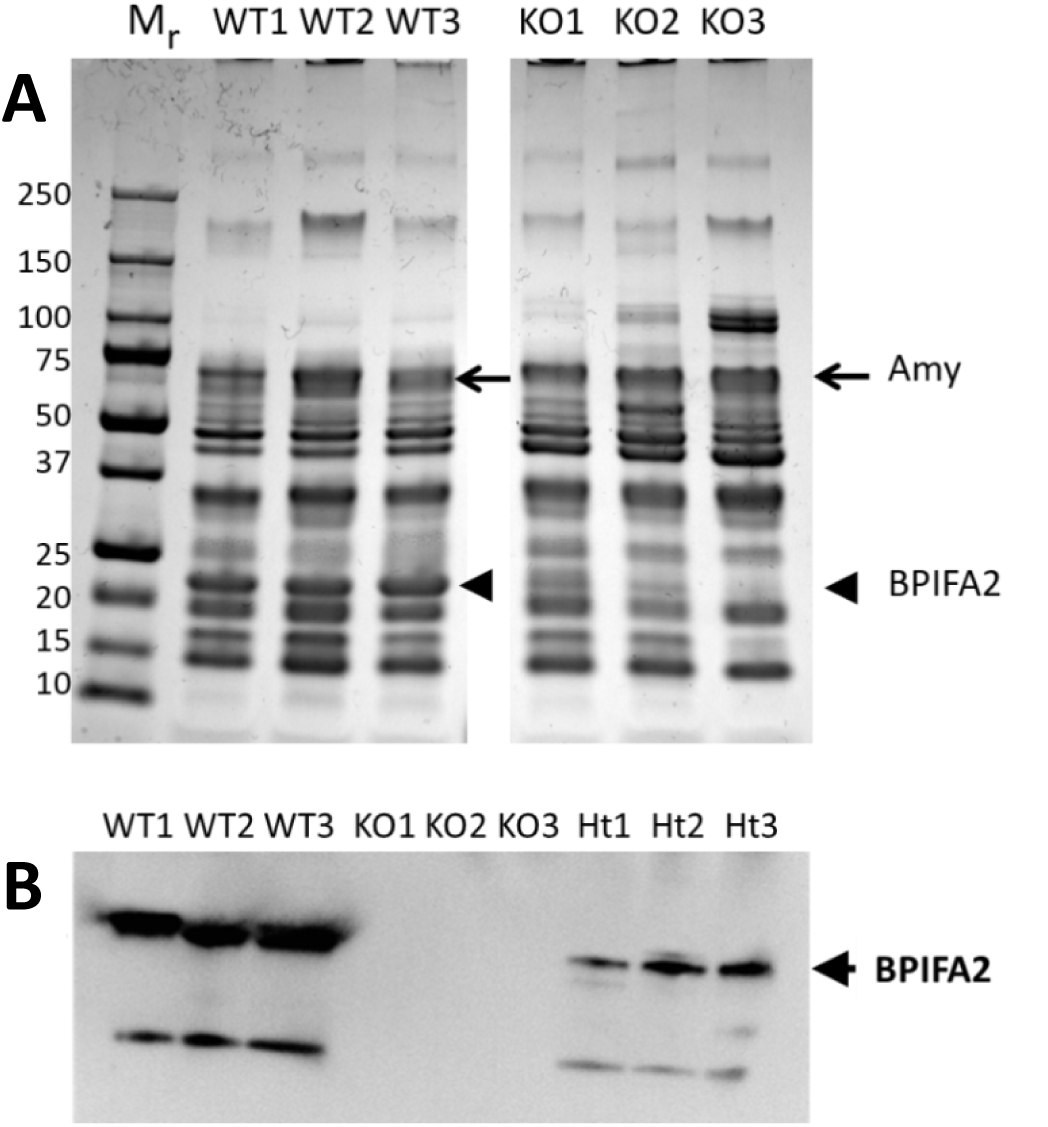
Analysis of mouse saliva. A. Saliva from three WT and three *Bpifa2* KO male mice was analyzed by SDS-PAGE. Molecular size standards (Mr, kD) are shown on the left and the expected positions of amylase (arrows) and BPIFA2 (arrowheads) are indicated. B. Detection of BPIFA2 in mouse saliva. Saliva from three WT, *Bpifa2* KO and heterozygous (Ht) mice, respectively, was analyzed with an antibody to mouse BPIFA2. The experiment was repeated with WT and *Bpifa2* KO saliva with similar results.

Since BPIFA2 is related to the surfactant proteins PLUNC (BPIFA1) (Bartlett et al., 2011) and latherin (BPIFA4) (Beeley et al., 1986; Vance et al., 2013) and the surfactant properties of saliva may be important for its function, we determined if BPIFA2 depletion affected the surfactant properties of mouse saliva. This was assessed by measuring the changes in contact angle of saliva droplets on a hydrophobic surface. Stimulated saliva collected from KO mice was less able to spread on a hydrophobic surface than WT saliva (Fig. 2A), resulting in a smaller decrease of the contact angle over time after drop placement (Fig. 2B).

**Fig. 2.**
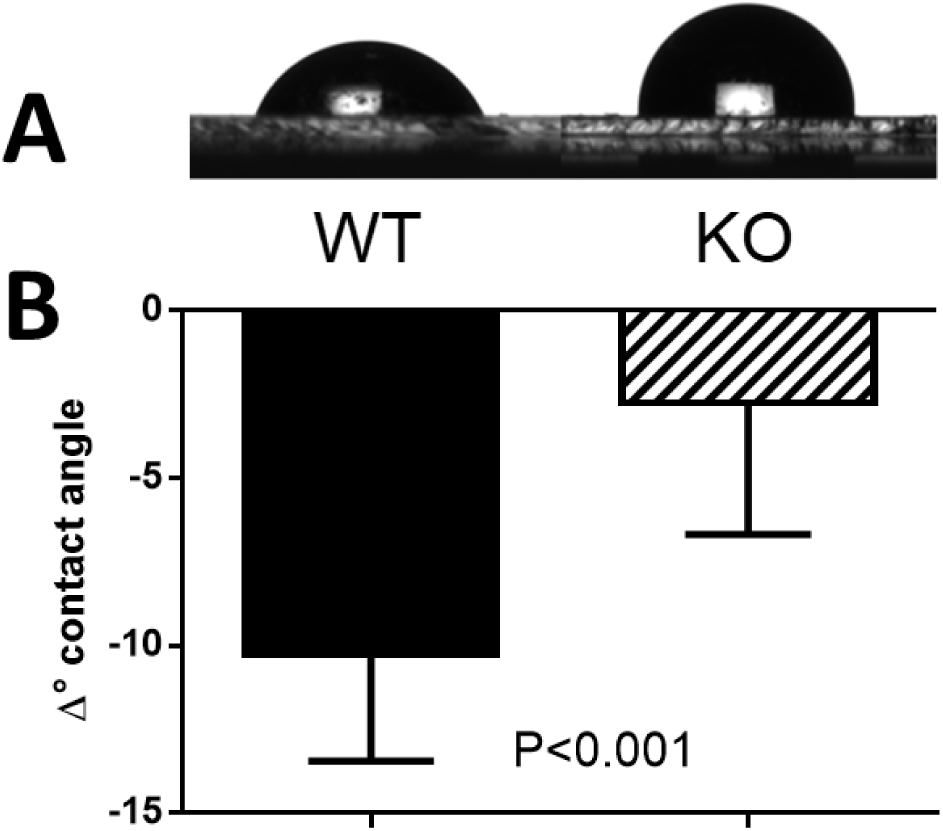
Saliva contact angles. **A**. drops of wild-type (WT) and knockout (KO) mouse saliva were deposited on a hydrophobic surface and photographed. **B**. The decrease in contact angle after placement of the droplet on a hydrophobic surface was determined by measuring the initial angle as well as that at 20s after placement of the droplet, and determining the difference. Samples from two mice per group were measured four times each. Student’s t-test, P-value is shown; N=8.

Another aspect of the surfactant property of saliva is its surface tension. The surface tension of mammalian saliva, including WT mouse saliva, is 45-50 mN/m (Fig. 3). In contrast, the surface tension of KO mouse saliva was found to be 70 ± 4 mN/m, which is close to that of water (Fig. 3).

**Fig. 3.**
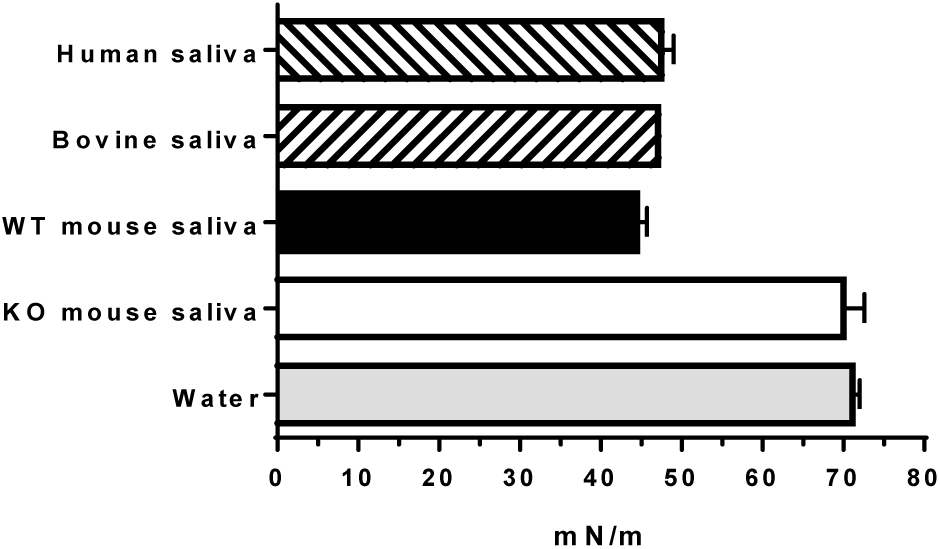
Surface tension of mammalian saliva. The surface tension of WT (black bar) and KO (open bar) mouse saliva was expressed as mean ± range; duplicate determinations from two mice per group. The published surface tensions of human saliva (Christersson et al., 2000; Freitas et al., 1993), bovine saliva (Reid and Huffman) and water (Gakhar et al., 2010; Reid and Huffman) are shown for comparison (mean ± range).

To determine if mice compensated for the absence of the surfactant activity of BPIFA2 by changes in saliva secretion, we determined the volume of pilocarpine-stimulated saliva collected from WT and KO mice. Saliva secretion was not altered in male mice but trended 15% lower in female KO mice compared to female WT mice. However, this difference did not quite reach statistical significance (Fig. 4). Thus, BPIFA2 depletion does not greatly affect saliva volume.

**Fig. 4.**
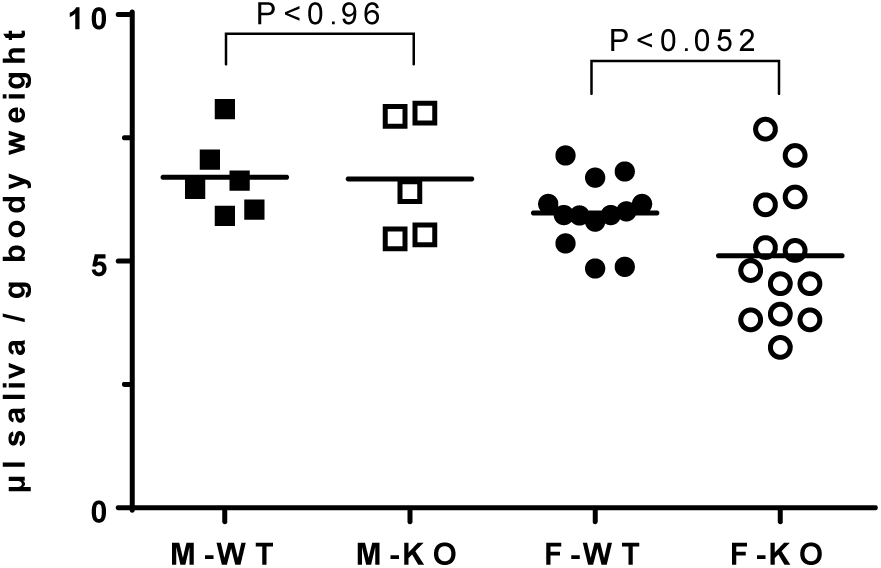
Saliva secretion. A. Pilocarpine-stimulated saliva was collected from male (M, squares) and female (F, circles), WT (closed symbols) and *Bpifa2* KO (open symbols) mice. Saliva volume was expressed relative to body weight (µl saliva / g body weight). Data from two (M) or four (F) independent experiments are shown with the group mean. WT and KO mice were compared by Student’s t-test for each sex (N=5-13).

To test if the watery saliva of *Bpifa2* KO mice affected their ability to form and swallow a food bolus, the consumption of dry powdered food was tested. The consumption of dry food did not differ between WT and KO mice of either sex (Fig. 5A). In addition, KO mice did not compensate for the lower surfactant activity of saliva by increased drinking. In fact, water consumption was reduced about 15% in KO mice compared to WT mice, although this only reached statistical significance in male mice (Fig. 5B).

**Fig. 5.**
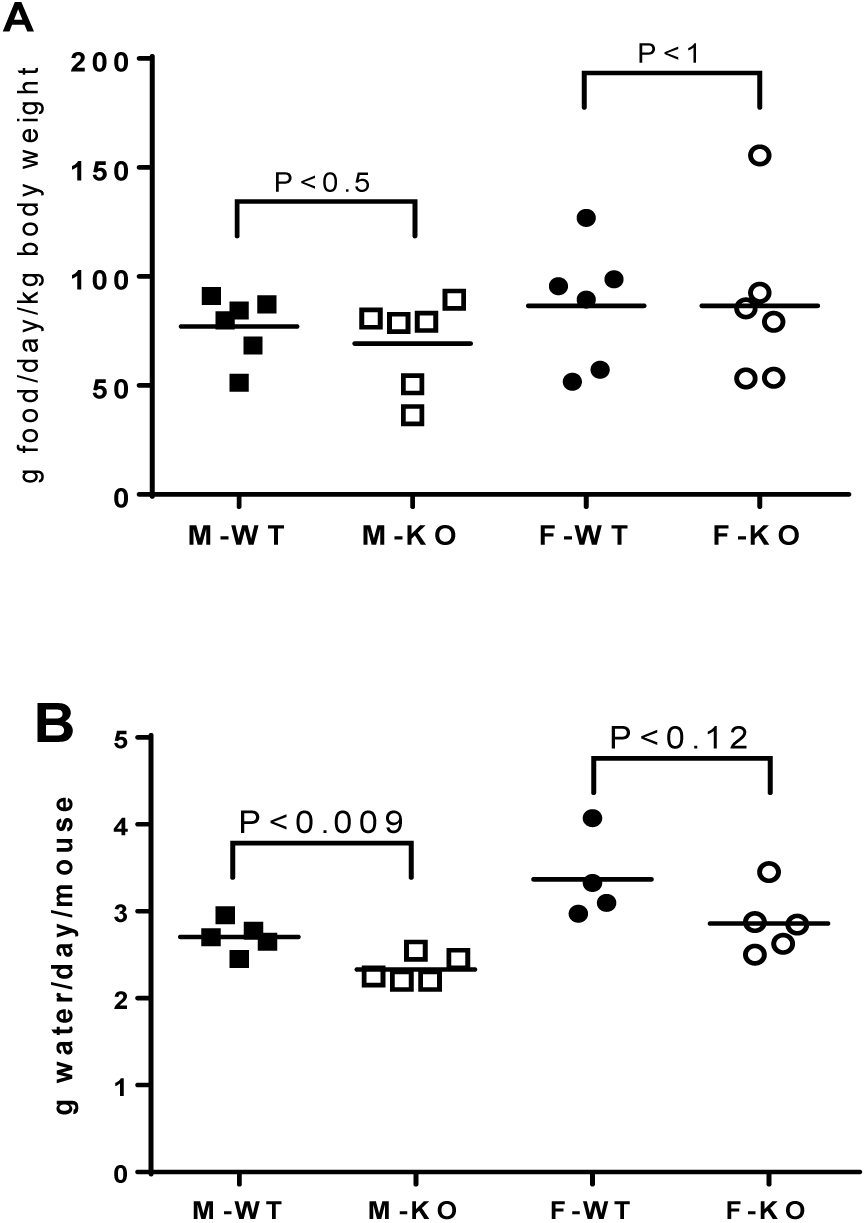
Consumption of food and water was determined for male (squares) and female (circles), WT (closed symbols) and KO mice (open symbols). A. Consumption of food (g/day/kg body weight). Data from three independent experiments were analyzed by unpaired Student’s t-test for each sex. P-values are shown, N=6 cages per group. B. Water consumption (g water/day/mouse) was determined for groups of 4 mice on 4-5 consecutive days. Data were analyzed by unpaired Student’s t-test for each sex. P-values are shown, N=4-5.

In addition to food intake (Amerongen and Veerman, 2002), saliva is also important for grooming and cooling in rodents (Roberts, 1988; Thiessen, 1988). We observed the grooming behavior of groups of WT and KO mice but did not note any differences in grooming or coat appearance. Moreover, WT and KO mice did not differ in their responses to an elevated ambient temperature. At 40-42°C, both groups coped by resting in an extended position (Roberts, 1988).

The somewhat reduced saliva secretion in female KO mice (Fig. 4) was reminiscent of reduced salivary flow in mouse models of Sjögren’s syndrome (Robinson et al., 1998b; Soyfoo et al., 2007). Since BPIFA2 has been implicated in the development of Sjögren’s syndrome-like symptoms in NOD mice (Robinson et al., 1998a; Robinson et al., 1998b), we tested if BPIFA2 depletion was associated with salivary gland inflammation. The number of inflammatory foci in submandibular glands did not differ between WT and KO mice of either sex (Fig. 7A). Linear regression indicated that the number of foci per tissue section increased by 0.5 per month as the mice aged, independent of sex (Fig. 7B) and genotype (Fig. 7C).

**Fig 7.**
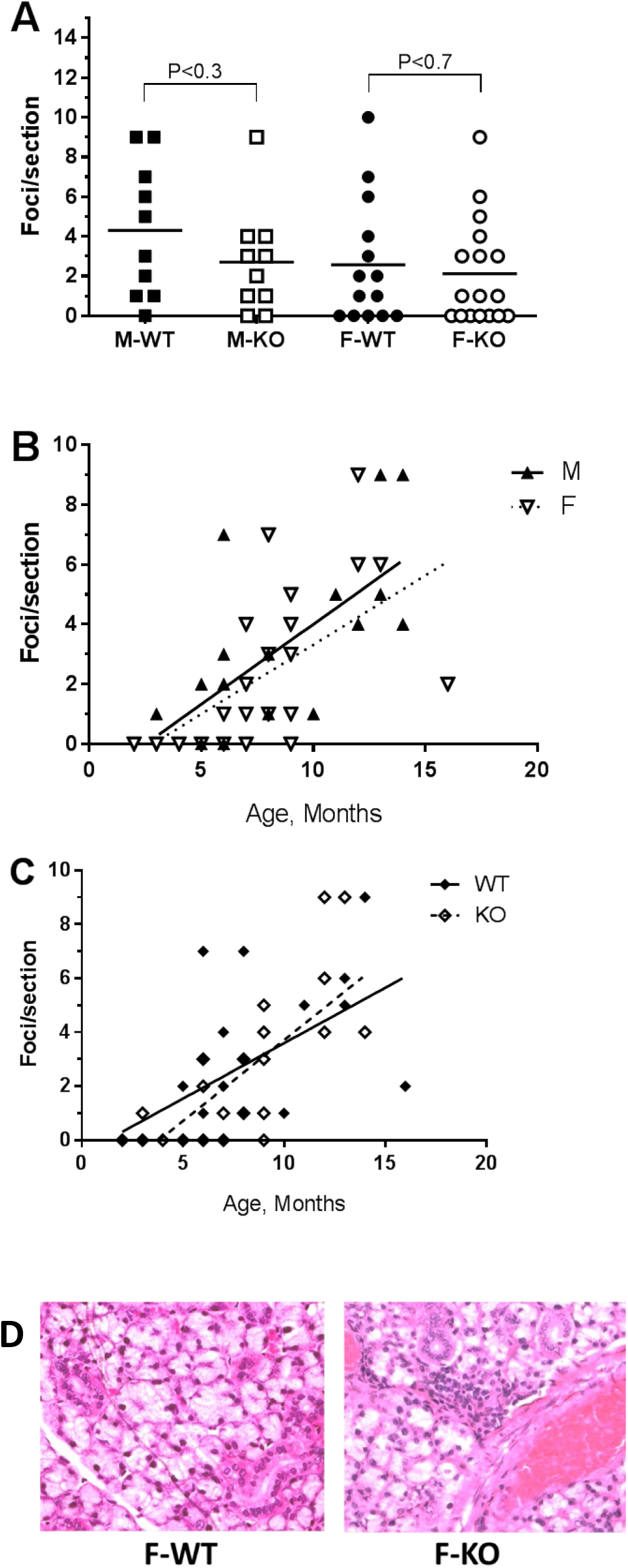
Inflammatory foci in submandibular gland. H&E stained tissue sections were analyzed by light microscopy and foci of inflammatory cells (>50 cells) counted per section. **A.** Foci per section for male (squares) and female (circles), WT (closed symbols) and KO mice (open symbols) glands. Data for individual mice are shown with mean. WT and KO mice of each sex were compared by unpaired, two-tailed Student’s t-test. P-values are shown; N = 10-17. **B.** Foci per section for WT and KO mice were combined for male (closed triangles) and female (open triangles) mice and expressed as a function of age. Linear regression slopes = 0.5; different from zero, P<0.003; N=19 (M) and P=0.0001; N=30 (F). **C.** Foci per section for male and female mice were combined for WT (closed diamonds) and KO (open diamonds) mice and expressed as a function of age. Linear regression: slope = 0.4 (WT) and 0.6 (KO); different from zero, P<0.007; N=22 (WT) and P<0.0001; N=27 (KO). **D.** H&E stained section of submandibular glands from 4 month old female WT and KO mice.

BPIFA2 is an LPS-binding protein (Abdolhosseini et al., 2012). To determine if BPIFA2 depletion alters the level of active LPS, LPS activity was determined in whole saliva and serum. Salivary LPS activity was reduced about 25% in KO mice, compared to WT mice (Fig. 8A), while serum LPS did not differ between WT and KO mice (Fig. 8B). Circulating LPS has been associated with the development of inflammatory loci in salivary glands (Wang et al., 2016). Thus, the lack of difference with genotype in levels of circulating LPS is consistent with the lack of differences in inflammatory loci observed in salivary gland sections (Fig. 7).

**Fig. 8:**
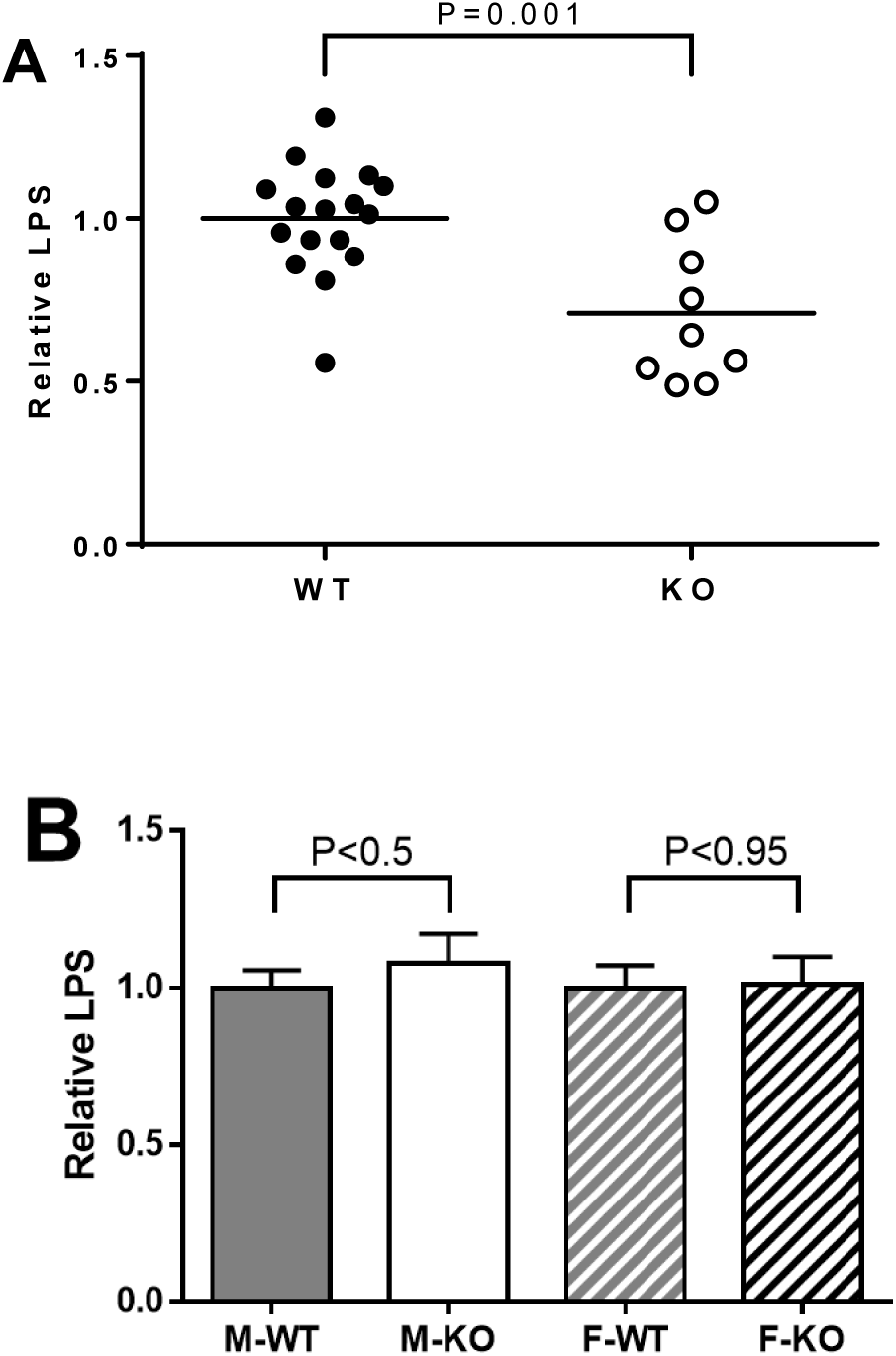
LPS analysis. **A.** LPS activity was determined in stimulated whole saliva from WT (closed symbols) and KO (open symbols) mice. The lines indicate the mean of each group. Data are from three independent experiments analyzed by unpaired, two-tailed Student’s t-test, N=9-17. P-value is shown. **B**. LPS activity was determined in blood plasma from mice that had been fasted 12h before blood collection. The data were normalized to the mean of the WT samples in each experiment and are shown as mean ± SEM. Data from three independent experiments were analyzed by unpaired, two-tailed Student’s t-test for each sex. P-values are shown (N=34)

Since WT and KO mice did not exhibit differences in circulating LPS, and glandular saliva can be considered sterile (Schroder et al., 2017), we reasoned that salivary LPS was derived from oral bacteria in anesthetized mice. When awake, mice also may ingest LPS from food and coprophagy (Soave and Brand, 1991). To directly test the effect of swallowed LPS on the GI tract, mice were fed LPS in their drinking water and the effect on intestinal motility was determined (Anitha et al., 2012). Fig. 9A shows that KO mice responded to ingested LPS by extruding 30% more fecal pellets than their WT colony mates. Stool frequency did not differ between WT and KO mice that were fed drinking water without added LPS. Similarly, the weight of the expelled feces did not differ significantly between WT and KO mice in the two treatment groups (Fig. 9B).

**Fig. 9:**
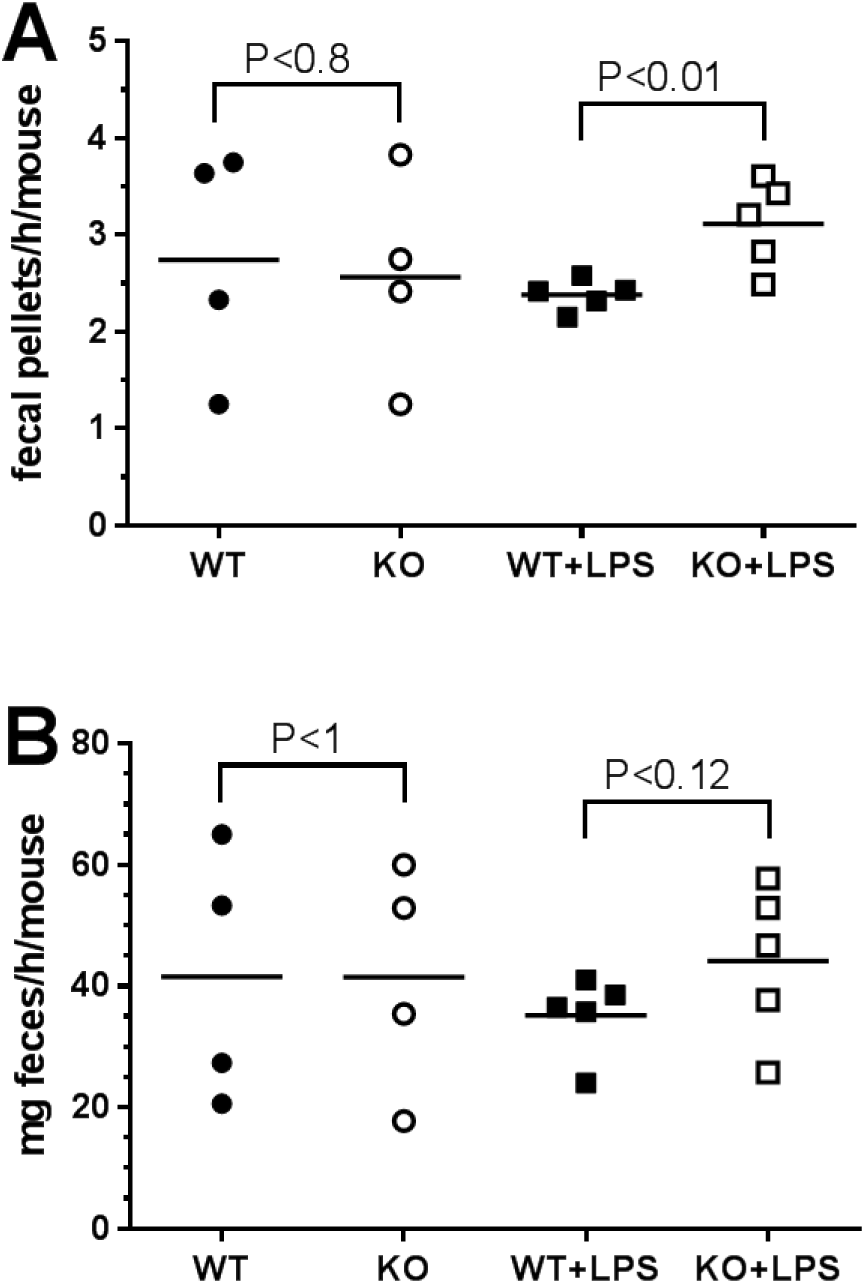
Effect of LPS on stool frequency and weight. Groups of male or female mice were fed drinking water without (circles) or with added LPS (squares) and the effect on stool frequency (A) (fecal pellets/h/mouse) or fecal mass (B) (mg feces/h/mouse) was determined in WT (closed symbols) and KO mice (open symbols). WT and KO mice in each treatment group were compared by paired, two-tailed Student’s t-test. P-values are shown; data are from 2-3 independent experiments, each with 1-2 cages/experimental group (N=4-5) and 2-4 mice/cage.

A preliminary histological analysis of the intestine, suggested morphological changes in KO mice (Gorr et al., 2015) but these changes were not confirmed by subsequent more detailed analysis (not shown). A sustained increased level of LPS activity in the intestine, while not affecting circulating LPS (Fig. 8B), could cause a low-grade endotoxemia, which has been associated with metabolic disorder and insulin resistance (Cani et al., 2007). WT and KO mice did not differ in their fasting glucose concentrations (Fig. 10A) while the KO mice of both sexes showed about 2-fold increased insulin levels after a short fast (<2h) (Fig. 10B).

**Fig. 10.**
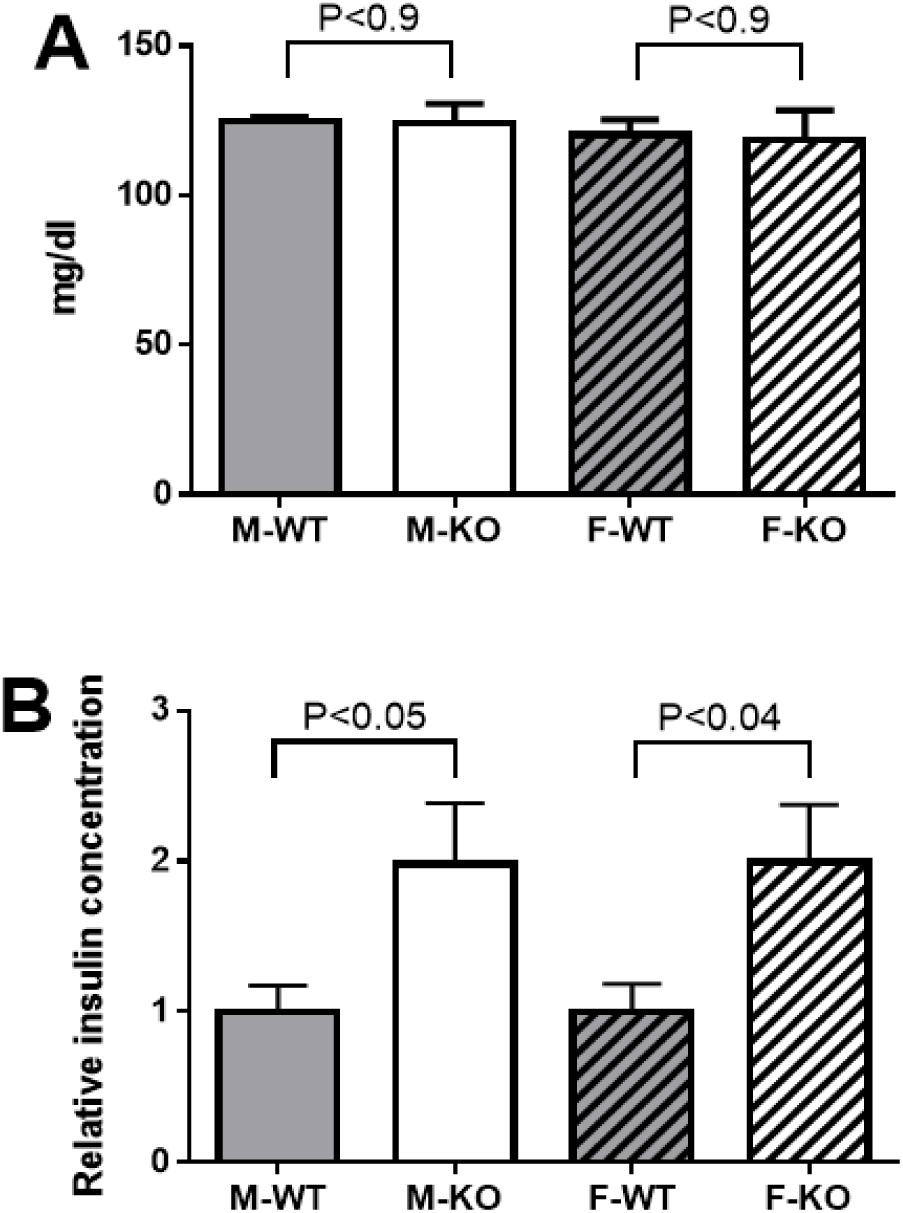
Glucose levels and insulin secretion. A. Blood glucose was determined in freshly collected blood from mice aged 6 months and expressed as mean ± SD, N=4. Similar results were obtained when the same mice were tested at ages 8-9 months (not shown). B. Relative insulin concentration in mouse serum. Data from three independent experiments were normalized to the mean of the WT group in each experiment for each sex and are shown as mean ± SEM. N=12-13. In each panel, WT and KO mice of each sex were analyzed by unpaired, two-tailed Student’s t-test.

While BPIFA2 depletion affects insulin levels, markers of lipid metabolism, including body fat, total cholesterol, triglycerides (IMPC, 2019) and fat droplets in the liver (not shown) were not altered by BPIFA2 depletion.

To test further the metabolic consequences of BPIFA2 depletion, targeted metabolomics was conducted using the Biocrates p180 platform for the analysis of acylcarnitines, amino acids, biogenic amines, glycerophospholipids, sphingolipids and sugars. The ratio of each metabolite detected in KO and WT mouse plasma was compared to the overall difference (P-value) between the two genotypes for male and female mice, respectively (Fig. 11). Only 4-5 of the 188 metabolites on the panel were not detected in these samples (Table 2).

**Fig. 11.**
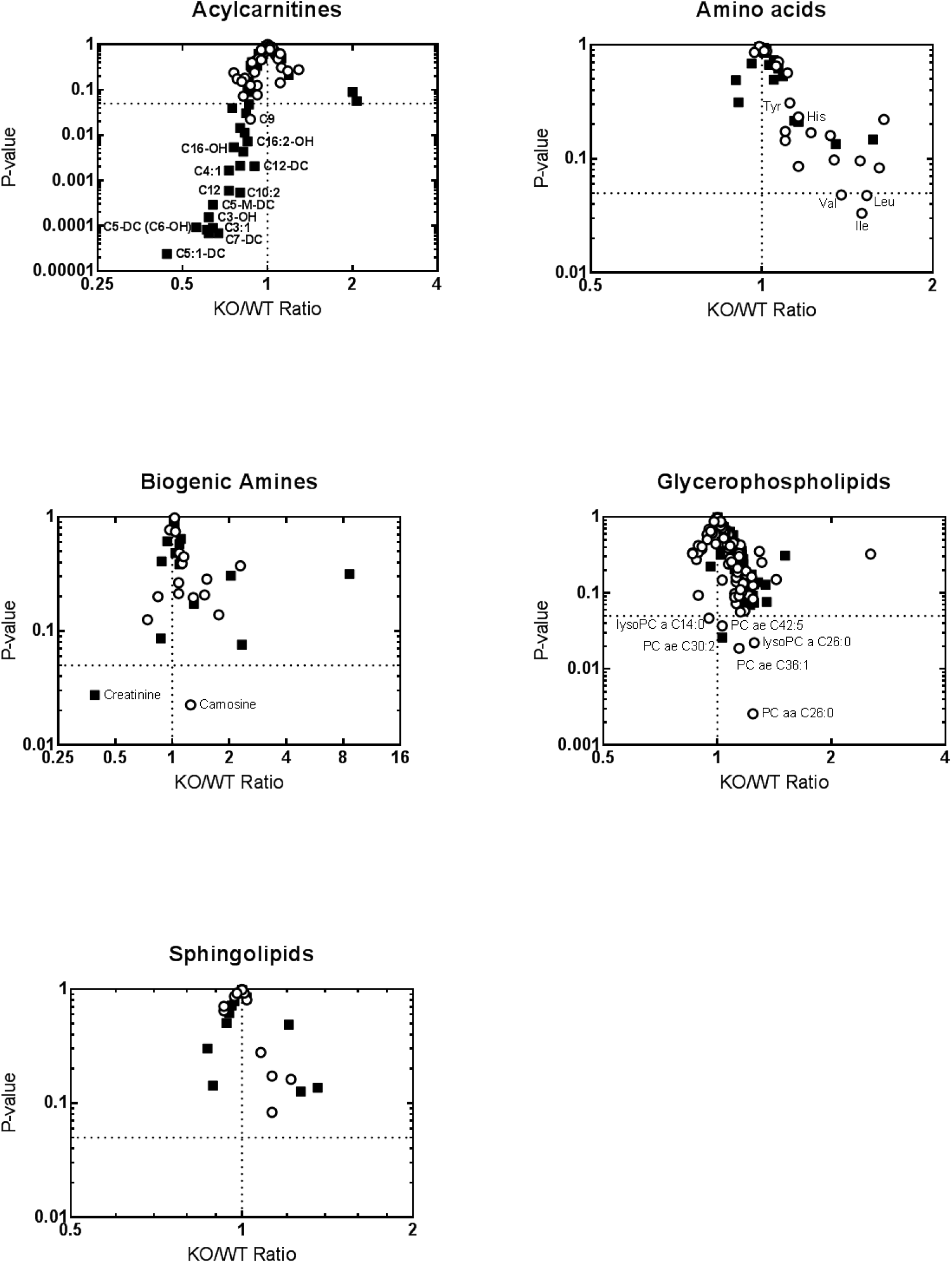
Targeted metabolomics of mouse plasma. Volcano plots show the KO/WT-ratio of the mean concentrations of 183 metabolites in male (filled squares) and female (open circles) mice.

**Table 2.** Targeted metabolomics of mouse plasma. Summary of metabolomic analyses shows the total number of metabolites analyzed in each group (On panel), the metabolites detected in mouse plasma (Detected) and the metabolites that showed a significant change (P<0.05) between WT and KO mice (KO different from WT) for male and female mice, respectively (N=8-10).

| Metabolites | On panel | Detected | KO different from WT |  |
| --- | --- | --- | --- | --- |
|  |  |  | Male | Female |
| Acylcarnitines | 40 | 40 | 21 | 1 |
| Amino acids | 21 | 21 | 0 | 3 |
| Biogenic amines | 21 | 16-17 | 1 | 1 |
| Glycerophospholipids | 90 | 90 | 1 | 5 |
| Sphingolipids | 15 | 15 | 0 | 0 |
| Sugars (hexose) | 1 | 1 | 0 | 0 |
| Total | 188 | 183-184 | 23 | 10 |

About half of the detected acylcarnitines were decreased in plasma from male KO mice (Fig. 11). Notably, C3 and C5 acylcarnitines showed the largest decreases in KO mice compared to WT mice. Interestingly, carnitine deficiency has been associated with incomplete fatty acid oxidation, which in turn may contribute to insulin resistance (Lopaschuk, 2016).

None of the 21 amino acids tested were significantly changed in the blood of male mice, while the branched chain amino acids Ile, Leu and Val, were upregulated by about 50% in female KO mice. A similar increase in branched chain amino acids has been associated with insulin-resistant obesity in both humans and rodent models (Lynch and Adams, 2014; Yoon, 2016).

In the biogenic amine group, creatinine was down-regulated in male KO mice while carnosine was slightly upregulated in female KO mice. Five glycerophospholipids were upregulated 25% or less in female KO mice, while male KO mice did not exhibit any notable changes in this group. Similarly, none of the sphingolipids were significantly changed in either male or female KO mice (Fig. 11, Table 2).

The concentrations of each metabolite in WT and KO mice for each sex were compared by Student’s t-test. N=10 male and 8 female mice for each genotype.

## Discussion

The BPI-fold protein family consists of proteins with a wide range of proposed functions that center on lipid binding and host-defense functions (Bingle and Gorr, 2004). In particular, BPIFA2 has previously been reported to bind lipid (Venkatesh et al., 2011) and HDL (Khovidhunkit et al., 2005), LPS (Abdolhosseini et al., 2012), bacteria (Gorr et al., 2011; Robinson et al., 1997) and yeast (Holmes et al., 2014). In addition, the protein shows antimicrobial activity (Khovidhunkit et al., 2005; Prokopovic et al., 2014) and may regulate epithelial sodium channels (Tarran et al., 2012). Consistent with earlier results for PLUNC (BPFA1) (Gakhar et al., 2010) and latherin (BPIFA4) (McDonald et al., 2009), we now find that BPIFA2 is also a salivary surfactant and that this salivary protein has systemic metabolic effects.

Surfactant proteins are relatively rare in nature (Cooper et al., 2017). While BPIFA2 is one of about eight highly abundant proteins in mouse saliva (Fig. 1), depletion of this single protein changes the physical properties of saliva from a typical mammalian saliva to those of water. Although BPIFA2 is abundant in mouse saliva, there are at least 8-10 major proteins remaining in KO saliva (Fig 1). Thus, we consider it unlikely that the effect of BPIFA2 depletion is a non-specific effect of general protein depletion. Indeed, the watery property of KO saliva is detected despite the presence of the remaining major salivary proteins, including amylase. A similar correlation between saliva “wateriness” (by total protein concentration) and BPIFA2 expression was recently reported in primates (Thamadilok et al., 2019). As an example, gorillas show 4-5-fold higher total protein concentration of their saliva than humans but BPIFA2 is the only one of 13 tested proteins that is upregulated in gorilla compared to human saliva. Mucins have traditionally been associated with saliva stickiness and coating of oral surfaces (Tabak, 2006).

However, neither Muc5B nor Muc7 were upregulated in gorilla saliva (Thamadilok et al., 2019). These results strengthen the suggestions that BPI-fold family proteins play a role in the wetting of mucosal surfaces. On the other hand, we did not determine any negative effects of the more watery saliva in KO mice. In particular, KO mice did not respond to the lack of BPIFA2 by increased saliva secretion or water consumption. In fact, these indicators were slightly reduced in female and male mice, respectively. Thus, it appears that the BPIFA2-depleted saliva can maintain oral physiology under the housing and feeding conditions tested, including the ability to consume dry powdered food. Similarly, BPIFA2 did not appear to play a role in grooming or temperature regulation. Consistent with the latter finding, it has been reported that ligation of parotid salivary ducts, the main source of BPIFA2 (PSP) in adult rat saliva (Mirels et al., 1998), does not alter temperature regulation in rats (Hainsworth, 1967).

BPIFA2 is an LPS-binding protein and we have previously identified a peptide in the BPIFA2 sequence that inhibits LPS activity (Abdolhosseini et al., 2012). The LPS content of BPIFA2-depleted saliva was about 25% lower than in WT saliva. Since the mice were anesthetized prior to saliva collection, this LPS level likely reflects reduced solubilization of LPS from oral bacteria. When the mice were directly challenged with ingested LPS, an increased rate of fecal pellet extrusion was observed in KO mice but not in WT mice. These results suggest that BPIFA2 affects the activity of ingested LPS. As such, BPIFA2 may play a protective function to prevent endotoxemia caused by ingested LPS. Interestingly, high levels of BPIFA2 expression have been reported in species that engage in coprophagy, including non-human primates (Thamadilok et al., 2019), dogs (Pasha et al., 2018; Torres et al., 2018), and rodents (Gao et al., 2018; Mirels and Ball, 1992; Venkatesh et al., 2004). Thus, it is possible that expression of this protein has evolved to facilitate nutrient extraction from feces or to protect against ingested LPS. Further studies with mice treated with oral antibiotics (Kesavalu et al., 2007) or prevented from coprophagy (Ebino et al., 1988) will be necessary to distinguish between oral and ingested sources of LPS.

Mild inflammation, triggered by bacterial LPS, has been linked with metabolic syndrome and insulin secretion (Cani et al., 2007). Correspondingly, gut-derived LPS has been associated with increased insulin secretion in both fasted and glucose-stimulated rats (Cornell, 1985) but increased insulin secretion was not associated with changes in plasma glucose concentrations in rats (Cornell, 1985). Thus, our finding that insulin secretion is increased in KO mice, without a change in glucose concentration, is consistent with the hypothesis that BPIFA2 depletion allows increased activity of ingested LPS in the GI tract where it may affect insulin secretion and cause insulin resistance (Cani et al., 2007). Obesity in humans has been associated with increased serum levels of insulin, branched chain amino acids and acylcarnitines, among other metabolites (Newgard et al., 2009). In addition to increased insulin levels in KO mice, metabolomic analysis detected increased levels of branched chain amino acids (Ile, Leu, Val) in female KO mice and a decrease in several acylcarnitines in male KO mice. On the other hand, serum lipid levels do not appear to be altered in KO mice, as determined by IMPC (IMPC, 2019). In a preliminary report, we noted increased hepatic lipid storage in KO mice (Gorr et al., 2015), but further analysis revealed that this was correlated with mouse age rather than BPIFA2 expression (not shown). The present study detected modest metabolic effects in KO mice that were fed standard, irradiated rodent chow. It is likely that enhanced effects would be detected under conditions that stress the metabolism, including high-fat diets (Ding et al., 2010), alcohol consumption (Tamai et al., 2002) or high-fructose diets (Mai and Yan, 2019). Overall, the results presented in this paper support the distal physiological function of a salivary protein. These findings reinforce the connection between oral biology and systemic disease. The BPIFA2 KO mouse opens up new avenues for research to establish the effect of this salivary protein on systemic health and disease.

## Acknowledgements

SRN current address: Department of Biochemistry & Molecular Biology, George Washington University, Washington D.C, USA.

We thank H. Hirt, University of Minnesota for assistance with initial assays and helpful discussions. S. Ruhl, University at Buffalo is thanked for helpful discussions of unpublished data. We thank staff at the University of Minnesota Research Animals Resource for assistance with colony maintenance and sample collection.

